# Exploring spatiotemporal changes in ecosystem service values and hotspots in Southwest of China, Mekong Region

**DOI:** 10.1101/2020.02.13.947317

**Authors:** Zhuoya Zhang, Junjie Yang, Xin Yang, Fuming Xie

**Author notes:** Corresponding author: Xin Yang, Communist Youth League Committee, Southwest Forestry University, No. 300, Bailong Road, Kunming, Yunnan, People’s Republic of China.

## Abstract

Xishuangbanna in the southwestern border of China is located in the upper reaches of the Mekong River between the Indo-China Peninsula and the East Asian continent. It is the largest tropical rain forest and monsoon forest in China. They have an irreplaceable ecological service function and are an important barrier to maintaining the ecological security of the Mekong River transboundary basin of the Lancang River. Based on three sets of remote sensing data (1996, 2003, 2010 and 2016), this study has made an exploration on spatio-temporal changes of Xishuangbanna, and also made an assessment on different ecosystem values of six types of ecosystem in Xishuangbanna based on related theories of ecological economics. Results showed that the values of Xishuangbanna ecosystem services in 1996, 2003, 2010, and 2016 were 70.4014billion yuan, 70.2115 billion yuan, 68.5129 billion yuan and 63.6098 billion yuan, respectively. The total value shows a continuous decreasing trend, reflecting the continuous decline in the ability of the Xishuangbanna ecosystem to provide services to humans. If divided by ecosystem types, forest and rubber were two types that have the greatest proportion of service values. In the perspective of service types, soil formation and protection account for the largest proportion, followed by gas regulation and biodiversity protection. Studying Xishuangbanna’s ecosystem service values is helpful to explore the sustainable development of resources and economy.

**Highlight:** This paper used coefficient of sensitivity to determine the dependence of ESV on the change of ecosystem value coefficient over time.

The total amount of ESV shows a continuous downward trend, indicating that the ecological environment of Xishuangbanna is still severe.

The temporal-spatial change of ESV was analyzed for two decades.

## Introduction

Ecosystem services (ES) refer to the various benefits that humans receive from ecosystems, including supply services, regulation services, support services and cultural services (Alcamo 2003, MEA 2005). The intensification of human activities has greatly accelerated the climate, environment and ecosystems of the planet. Urban ecosystems are increasingly threatened by urban population growth, urban land use expansion(Storkey, Döring et al. 2015, Leverkus and Castro 2017, Yang, Guan et al. 2018, Kuriqi, Pinheiro et al. 2019), and socio-economic activities(Mensah, Veldtman et al. 2017, Song and Deng 2017), which affects the value of ecosystem services and ultimately the sustainable development of human society(Costanza, d’Arge et al. 1997, Sun, Liu et al. 2016). Accurate assessment of ecosystem services value (ESV) is essential for urban construction planning and the improvement and restoration of urban ecosystems(Cui, Xiao et al. 2017), so it has received increasing attention from the research community(Costanza, Chichakly et al. 2014, Costanza, De Groot et al. 2014, Leverkus and Castro 2017, Ossola and Hopton 2017).

As Costanza et al proposed the value of ecosystem services, especially after the Millennium Ecosystem Assessment, the impact of human activities on ecosystems has become an important research direction(Costanza, d’Arge et al. 1997, Robert, Costanza et al. 2005). Many scholars have emphasized that human activities are an important driving force for changing ecosystem services(Danz, Niemi et al. 2007, Robards, Schoon et al. 2011). The true economic value of ecosystem services depends on the interaction between the supply of ecosystems and the needs of society(Braat and de Groot 2012) (Robert, Costanza et al. 2005). Monetary valuation of the importance of ecosystem services to society can serve as a powerful and essential communication tool to provide a basis for better and more balanced decisions(De Groot, Brander et al. 2012). Scholars have found that basic income transfers can be a convenient way to determine ESVs globally and nationally, assuming a constant unit value per hectare for a given ecosystem type multiplied by the area of each type to arrive at a total (Costanza, De Groot et al. 2014).

The value of ecosystem services not only reflects the functions of ecosystem services, but also reflects the significance of human ecological environment and the demand for ecosystem services. Similarly, many scholars have analyzed the spatio-temporal changes of ESV, and their responses to land-use changes(Fengqin, Yan et al. 2016, Kindu, Schneider et al. 2016, Fei, Shuwen et al. 2018) or other human activities(Camacho-Valdez, Ruiz-Luna et al. 2014). Although it is important to have a better understanding of the temporal changes of ESVs, there is an increasing focus on determining the spatial changes of ESVs by identifying “hot spots” on ecosystem services(Li, Fang et al. 2016). These spatial studies can provide a range of useful tools that can effectively integrate ecosystem services into planned or current conservation plans(Naidoo, Balmford et al. 2008)], assess the effectiveness of implementing ecological policies, and prioritize the management of ecosystem services field(Egoh, Reyers et al. 2011, Li, Fang et al. 2016). This information is particularly important for modifying current ecological protection plans and policies in a more beneficial and targeted manner. For example, by linking the spatial changes in wetland area with the provision of ecosystem services and economic value, some scholars have analyzed the spatiotemporal changes in the service value of coastal landscapes in Southern Sinaloa (Mexico) (Camacho-Valdez, Ruiz-Luna et al. 2014). Bottalico [28] evaluated the spatial distribution of Molise ESV by developing a spatial explicit method(Bottalico, Pesola et al. 2016). Li et al. verified the spatiotemporal changes of ESV and its hot and cold issues in China(Li, Fang et al. 2016).However, ESVs are less well-characterized by hotspots that change over a specific spatial range, especially globally and nationally.

With the change of policies in Xishuangbanna, economic development and the advancement of urbanization, the transformation and occupation of land resources has intensified, resulting in a periodical sharp change in land use types and areas. This change has a great impact on ESV in Xishuangbanna. In this paper, we investigated the spatiotemporal changes of ESV and identified hotspots and hotspots of ESV changes. The analysis was based on Xishuangbanna four-phase remote sensing image data from 1996 to 2016. In this context, studying the impact of ecosystem services in Xishuangbanna from the perspective of land use change in Xishuangbanna can clarify the state of the ecosystem under the “stress-state-response” in Xishuangbanna. This is of great significance for adjusting and optimizing the land use pattern in Xishuangbanna, promoting the coordination of sustainable development in Xishuangbanna, and protecting the stability of cross-border ecological security.

## 1. Methods

### 1.1 Study area

Xishuangbanna is located at 21 ° 10′-22 ° 40 ′ north latitude and 99 ° 55′-101 ° 50 east longitude. It is on the tropical edge south of the Tropic of Cancer. The land area is 19,124.50 square kilometers. It is bordered by Puer City in the northeast and northwest, Laos in the southeast, and Myanmar in the southwest. The border line is 966.3 kilometers long. The highest altitude is 2429 meters and the lowest altitude is 477 meters. The whole state has jurisdiction over one city and two counties, Jinghong City, Menghai County, and Mengla County. The climate of Xishuangbanna is warm and moist all year round. There are no four seasons except the dry and wet seasons. The dry season runs from November to April of the year and the wet season runs from May to October.

### 1.2 Data soures and processing

The social and economic data of Xishuangbanna include the publicly released relevant data such as the website of the National Bureau of Statistics of the People’s Republic of China and the statistical yearbook of Xishuangbanna. The remote sensing image data include: availability images using vegetation-free cloud-free datasets, which consist of four Landsat TM / TM + / OLI remote sensing images from March 1996 to April 1996, 2003, 2010, and April 2016, respectively.

Based on field surveys, combined with remote sensing data spectral information, we referenced the second-level survey data of forest resources in 2006 and 2016 to construct a remote sensing interpretation mark for land use in Xishuangbanna. Support Vector Machine (SVM) supervised classification method was used for The land use type data were interpreted and the land use types in the fourth phase of the study area were obtained. After the four-phase image classification, the total accuracy Kappa coefficients are 85.9%, 86.7%, 89.9%, and 93.5%, which meet the research needs.

### 1.3 Calculation of ESV in Xishuangbanna

#### 1.3.1 Ecosystem classification

According to China’s latest classification standard for land use status (GB / T21010-2007), and combined with the actual local conditions in Xishuangbanna, the ecosystem in Xishuangbanna is divided into forest ecosystem, rubber ecosystem, tea garden ecosystem, farmland ecosystem, watershed ecosystem, and construction land ecosystem.

#### 1.3.2 Evaluation method of ESV in Xishuangbanna

##### (1) Modification of Xishuangbanna’ESV Equivalent Factor

In this study, the tea garden ecosystem was taken as the average value of woodland and grassland(Xiao-qing, Ze-xian et al.). The forest ecosystem equivalent factor in Xishuangbanna was revised to 1.96 times the national average(Xiaosai, Yongming et al. 2015), and the rubber ecosystem equivalent factor was revised to 1.60 times the national average(Yuanfan, Qingzhong et al. 2010). Based on the above amendments, the Xishuangbanna ESV Equivalent Scale was obtained.

##### (2) Functional value of food production per unit area of cultivated land ecosystem in Xishuangbanna

In order to eliminate the influence of crop price fluctuations on the total value in each year and region, the basic data were selected from the sown area, total output and average price of three main crops (rice, upland rice, and corn) in 1997 in Xishuangbanna. According to Xie Gaodi’s research, the economic value of a standard unit ecosystem service value equivalent factor is equal to 1/7 of the average market value of a single grain(Gaodi, Chunxia et al. 2003, Xiaosai, Yongming et al. 2015). The calculation is as follows:

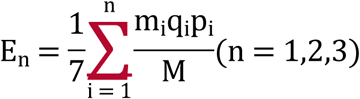

Among them, E_n_ is the economic value (yuan / hm^2^) of providing food production and service functions for a unit of cultivated land ecosystem. i is the type of crop. p_i_ is the price of icrops (yuan / kg). q_i_ is the yield of i food crops (kg / hm^2^). m_i_ is the area of i food crops (hm^2^). M is the total area of n food crops (hm^2^).

##### (3) Calculation method of Xishuangbanna ‘ESV

We used the ESV formula of Costanza et al. to calculate the ESV of Xishuangbanna (Robert, Costanza et al.). In order to facilitate the analysis of the changes in the value of ecosystem services in Xishuangbanna and to compare the value of ecosystem services in different regions, this study adds two indicators to the calculation method of ecosystem service values, the Ecosystem Services Contribution Rate (ESCR) and Ecosystem Services Value per unit (UESV) (Tianhai, field et al. 2018). The value of the ecosystem per unit area is to reduce the difference in the size of different ecosystems and to facilitate comparison. The calculation is as follows:

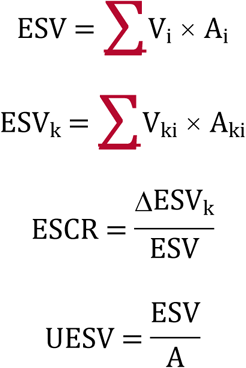

Among them, ESV is the total value of ecosystem services in the study area. V_i_ is the ecosystem service value coefficient of ecosystem type i per unit area. A_i_ is the area of ecosystem type in the study area. ESV_k_ is the value of the k-th service of the ecosystem. V_ki_ is the k-th ecosystem service value coefficient of the i-th ecosystem type. ESCR is the change in the value of a single ecosystem serviceΔESV_k_ to the total value of ecosystem services. UESV is the ratio of total ecosystem service value ESV to total area A.

##### (4) Calculation method of Coefficients of Sensitivity(CS)

This study applied CS to measure the representativeness of individual ecosystem services to each ecosystem type and the accuracy of the value of ecosystem services per unit area (Yao, Rusong et al. 2012). CS is the change in the ecosystem service value coefficient V_i_ of the ecosystem type i per unit area caused by a 1% change in the ESV of the total ecosystem service value in the study area. CS> 1, which indicates that ESV is elastic to V_i_ and the accuracy of V_i_ is low. CS <1, it indicates that ESV is inelastic to V_i_, which indicates that V_i_is more accurate and ESV estimation is accurate. The value of the ecosystem service value of each ecosystem type is increased or decreased by 50%, and the change of the elasticity coefficient of the total value of ecosystem services is analyzed (Xiao-qing, Ze-xian et al., Yao, Rusong et al. 2012). The calculation is as follows:

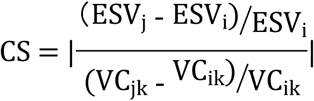

Among them, ESV is the total value of ecosystem services in the study area. VC is the coefficient of ecosystem service value per unit area. A_i_ is the area of ecosystem type in the study area. k is the kth service of the ecosystem. i and j are the adjusted values of initial value and ecological value coefficient.

## 2 Results

### 2.1 Changes in total values of ecosystem services in Xishuangbanna from 1996 to 2016

#### 2.1.1 ESV of different ecosystem types in Xishuangbanna from 1996 to 2016

According to calculations, the values of different ecosystem services from 1996 to 2016 are shown in Table 1.

Among the ESV of different ecosystem types in Xishuangbanna, the ecological value coefficients of forest ecosystems and rubber ecosystems are high. Changes in the area of such ecosystems will greatly affect the changes in ecosystem service values in Xishuangbanna.

**Table 1.**
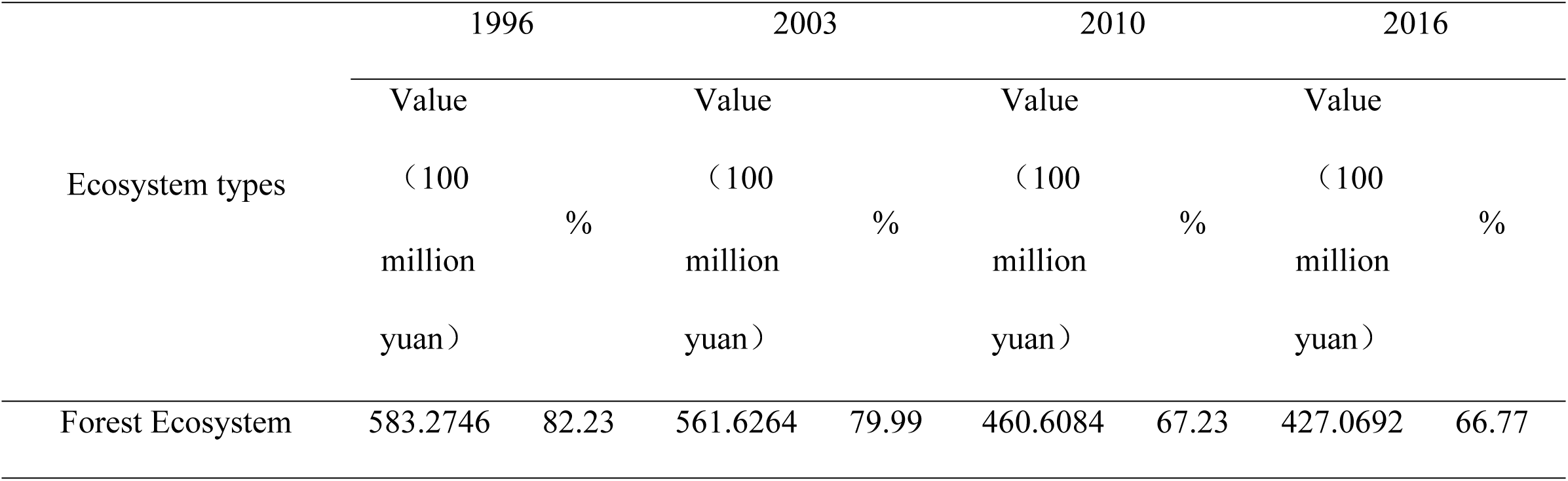
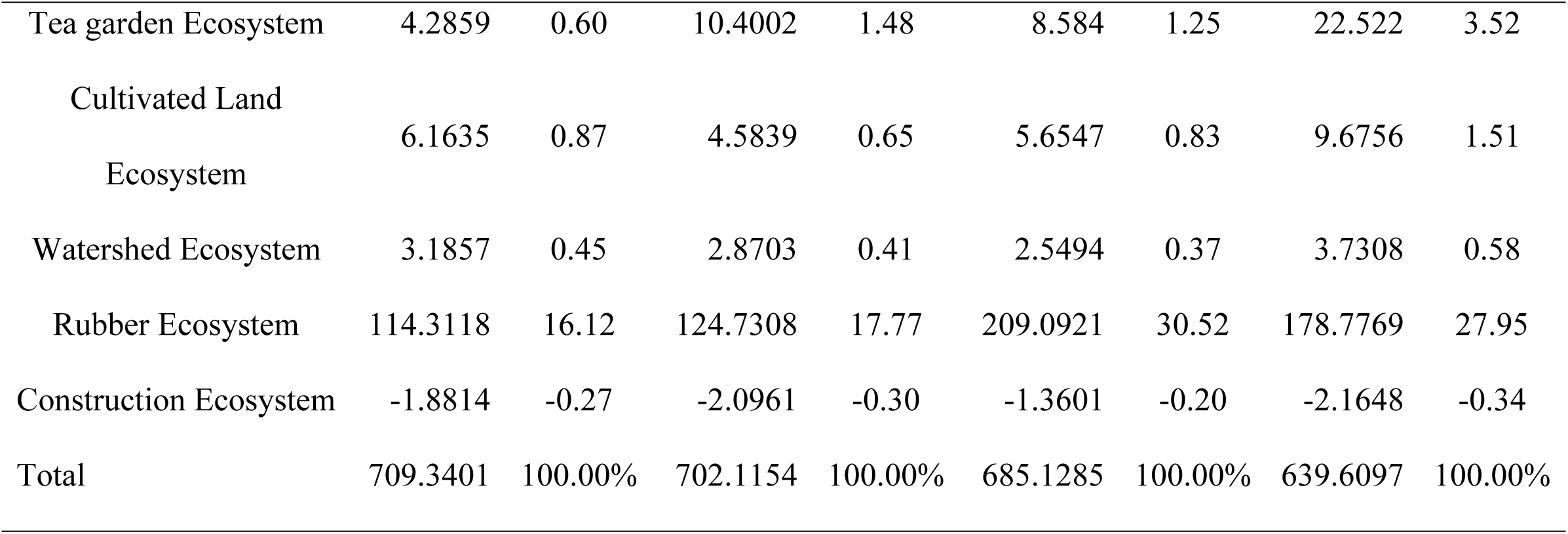
Ecosystem service values of each ecosystem type in Xishuangbanna from 1996 to 2016.

|  | 1996 |  | 2003 |  | 2010 |  | 2016 |  |
| --- | --- | --- | --- | --- | --- | --- | --- | --- |
|  | Value |  | Value |  | Value |  | Value |  |
| Ecosystem types | (100 | % | (100 | % | (100 | % | (100 | % |
|  | million |  | million |  | million |  | million |  |
|  | yuan) |  | yuan) |  | yuan) |  | yuan) |  |
| Forest Ecosystem | 583.2746 | 82.23 | 561.6264 | 79.99 | 460.6084 | 67.23 | 427.0692 | 66.77 |
| Tea garden Ecosystem | 4.2859 | 0.60 | 10.4002 | 1.48 | 8.584 | 1.25 | 22.522 | 3.52 |
| Cultivated Land<br>Ecosystem | 6.1635 | 0.87 | 4.5839 | 0.65 | 5.6547 | 0.83 | 9.6756 | 1.51 |
| Watershed Ecosystem | 3.1857 | 0.45 | 2.8703 | 0.41 | 2.5494 | 0.37 | 3.7308 | 0.58 |
| Rubber Ecosystem | 114.3118 | 16.12 | 124.7308 | 17.77 | 209.0921 | 30.52 | 178.7769 | 27.95 |
| Construction Ecosystem | -1.8814 | -0.27 | -2.0961 | -0.30 | -1.3601 | -0.20 | -2.1648 | -0.34 |
| Total | 709.3401 | 100.00% | 702.1154 | 100.00% | 685.1285 | 100.00% | 639.6097 | 100.00% |

From Table 1, it can be concluded that the ESV of Xishuangbanna in 1996, 2003, 2010, and 2016 showed a continuous decrease, which reflects that the ability of Xishuangbanna ecosystem to provide services to human beings has been continuously decreasing. The values of Xishuangbanna ecosystem services in 1996, 2003, 2010, and 2016 were 70.4014billion yuan, 70.2115 billion yuan, 68.5129 billion yuan and 63.6098 billion yuan, respectively. The total value shows a continuous decreasing trend, reflecting the continuous decline in the ability of the Xishuangbanna ecosystem to provide services to humans.

Among them, in terms of the proportion of ecosystem service value, the forest land ecosystem has the highest proportion, which is the most important component of the ESV in Xishuangbanna (66.77% −82.23%), followed by rubber ecosystem (16.12% −30.52%). Xishuangbanna is an area with high forest coverage, and the ecological environment is generally well preserved. However, with the development of human society and economy, the ecosystem services in Xishuangbanna show a continuous decline. In the case where the forest land and rubber ecosystem in Xishuangbanna account for a large proportion, the ESV of the forest land ecosystem and rubber ecosystem accounts for 94.71% −98.35% of the total value. It can also be seen that woodland and rubber ecosystems dominate the ecosystem services in Xishuangbanna.

The construction land ecosystem is the lowest and has a negative value, and the service value of the construction land ecosystem continues to show negative growth, from −6.6847 billion yuan in 1996 to −7.6916 billion yuan, which is also in line with the reality of the increasing area of construction land.

According to research, the ESV of rubber in Xishuangbanna is far less than that of tropical rain forests and secondary vegetation (67.149 million yuan / (hm^2^.a)), the proportion of which is about 56.74%. The ecological service value of evergreen coniferous forest (13,315 yuan / (hm^2^.a)) is large(Gaodi, Chunxia et al. 2003). The service value of the rubber forest ecosystem in Xishuangbanna is far greater than the average value of tropical forests in China (16.056 million yuan / (hm^2^.a)) (Yuanzhao 2003). The ecological and economic value of rubber forests is only about half of the value of tropical rain forests and secondary vegetation. Cutting down tropical rain forests and planting rubber forests is not desirable from the total regional ecological and economic value, which is not conducive to regional natural asset appreciation and local sustainable development(Tiyuan, Jiayong et al. 2009).

The direct economic output of rubber forest is much larger than that of tropical rain forest. This is the source of huge economic pressure on tropical rain forest protection. It is recommended to adopt environmental and economic policies such as ecological compensation to comprehensively protect tropical rain forest(Tiyuan, Jiayong et al. 2009).

#### 2.1.2 Changes in ESV of different ecosystem types in Xishuangbanna from 1996 to 2016

The total reduction of ESV from 2010 to 2016 (16.17153 billion yuan) was 630.19% of the total reduction of ESV from 1996 to 2003, which is the time period when ESV changes the most. During 2010-2016, the rapid development of urbanization, the continuous decline in rubber prices, the rise in the prices of tea and crops, and the continuous advancement of the country’s policy of “returning rubber to forests” have made the value of the Xishuangbanna ecosystem the most significant. Total reduction in ESV from 2003 to 2010 (6.03301 billion yuan). The period from 1996 to 2003 was the smallest total reduction in ESV (2.556614 billion yuan).

The ESV of forest ecosystems continues to decline. The change of forest ecosystem ESV is the most dramatic change among all ecosystem types, from 82.23% of the total ESV in 1996 to 66.77% of the total ESV in 2016. The period when the ESV of forestland ecosystem changed the most was from 2003 to 2010. During this period, especially from 2003 to 2007, the continuous increase of rubber prices in the world has caused a large number of forest land ecosystems to be replaced by rubber ecosystems. The rubber ecosystem ESV has increased sharply and the forest land ecosystem ESV has decreased sharply. The total reduction in ESV of forest land ecosystem from 2003 to 2010 (35.87842 billion yuan) is 64.67% of the total reduction from 1996 to 2016 (55.49324 billion yuan).

The ESV of the tea garden ecosystem decreased first, then increased sharply, and increased overall. From 2003 to 2010, the ESV of the tea garden ecosystem decreased. During this period, the tea price in Xishuangbanna was low, and rubber was the main source of the local economy. From 2010 to 2016, the tea market in Xishuangbanna was opened. The prices of tea, especially those of several famous tea mountains, rose sharply, and the ESV (4.995 billion yuan) of the tea garden ecosystem increased sharply.

The ESV of cultivated land ecosystem showed a trend of first decline, then slow increase, and overall growth. The ESV of cultivated land ecosystem decreased by 56106 million yuan from 1996 to 2003, and continued to increase from 2003 to 2010 and from 2010 to 2016. Among them, the biggest change was from 2010 to 2016. The total increase in ESV of forest land ecosystem (1.42819 billion yuan) from 2010 to 2016 was 114.49% of the total increase of ESV of forest land ecosystem (1.224746 billion yuan) from 1996 to 2016.

The ESV of water ecosystems are decreasing first, then increasing, and generally increasing slowly.

### 2.2 Changes of total values in single ecosystem service in Xishuangbanna from 1996 to 2016

#### 2.2.1 ESV in the individual ecosystem services in Xishuangbanna from 1996 to 2016

It can be seen from Table 2 that among the various ESVs in Xishuangbanna, soil formation and protection account for the largest proportion (17.86% −17.93%), followed by air regulation (accounting for 15.80% −15.92%) and biodiversity protection (accounting for 14.86% −14.91%), water conservation (accounting for 14.48% −14.57%), climate regulation (accounting for 12.34% −12.35%), raw material production (accounting for 11.61% −11.78%), and food production accounting for the smallest proportion (accounting for Than 0.56% −0.70%).

**Table 2.**
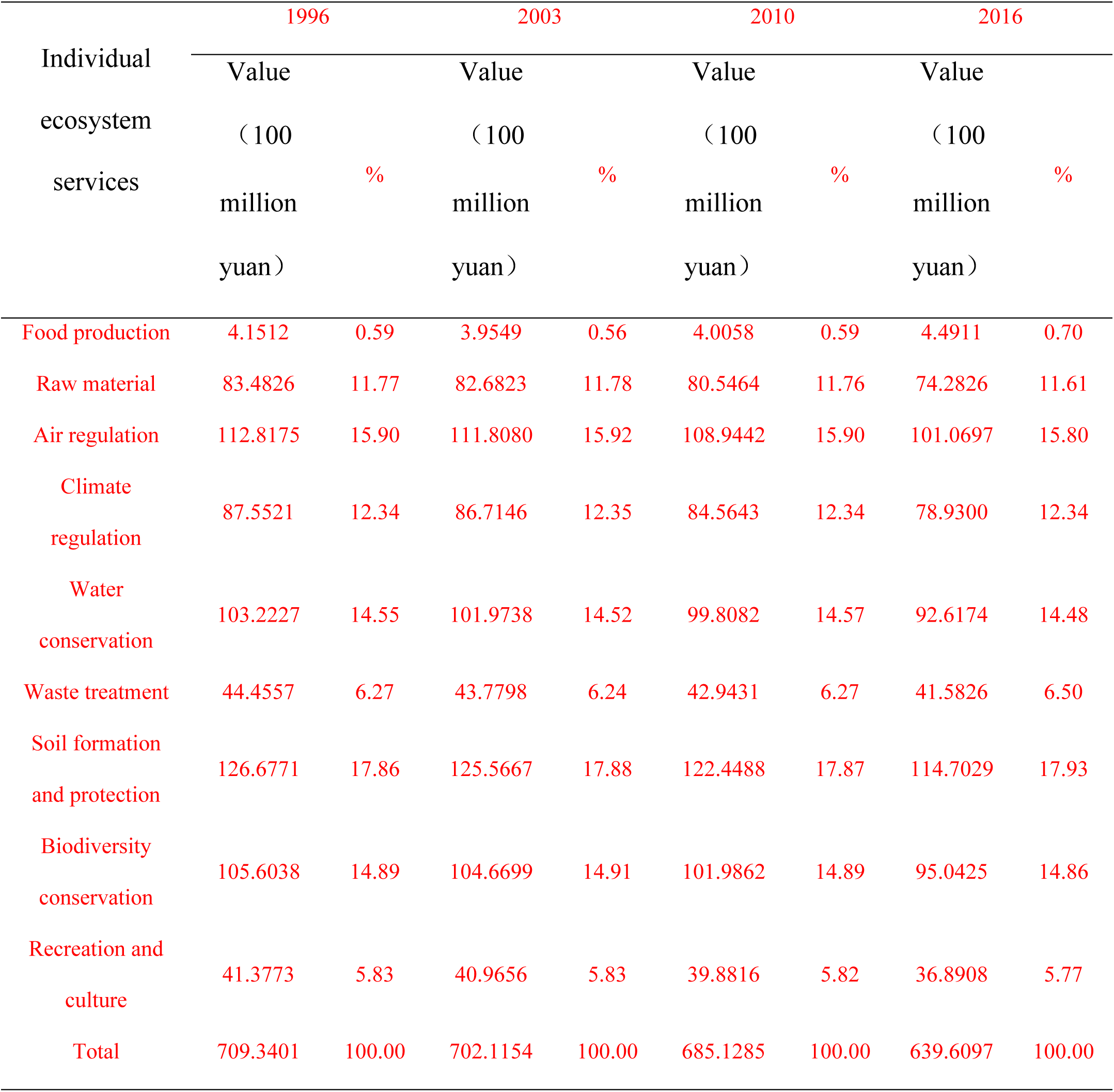
Evaluation of ecosystem service of different types in Xishuangbanna from 1996 to 2016.

|  | 1996 |  | 2003 |  | 2010 |  | 2016 |  |
| --- | --- | --- | --- | --- | --- | --- | --- | --- |
| Individual ecosystem services | Value<br>(100 million yuan) | % | Value<br>(100 million yuan) | % | Value<br>(100 million yuan) | % | Value<br>(100 million yuan) | % |
| Food production | 4.1512 | 0.59 | 3.9549 | 0.56 | 4.0058 | 0.59 | 4.4911 | 0.70 |
| Raw material | 83.4826 | 11.77 | 82.6823 | 11.78 | 80.5464 | 11.76 | 74.2826 | 11.61 |
| Air regulation | 112.8175 | 15.90 | 111.8080 | 15.92 | 108.9442 | 15.90 | 101.0697 | 15.80 |
| Climate regulation | 87.5521 | 12.34 | 86.7146 | 12.35 | 84.5643 | 12.34 | 78.9300 | 12.34 |
| Water conservation | 103.2227 | 14.55 | 101.9738 | 14.52 | 99.8082 | 14.57 | 92.6174 | 14.48 |
| Waste treatment | 44.4557 | 6.27 | 43.7798 | 6.24 | 42.9431 | 6.27 | 41.5826 | 6.50 |
| Soil formation and protection | 126.6771 | 17.86 | 125.5667 | 17.88 | 122.4488 | 17.87 | 114.7029 | 17.93 |
| Biodiversity conservation | 105.6038 | 14.89 | 104.6699 | 14.91 | 101.9862 | 14.89 | 95.0425 | 14.86 |
| Recreation and culture | 41.3773 | 5.83 | 40.9656 | 5.83 | 39.8816 | 5.82 | 36.8908 | 5.77 |
| Total | 709.3401 | 100.00 | 702.1154 | 100.00 | 685.1285 | 100.00 | 639.6097 | 100.00 |

Compared with urban areas with low forest coverage, this ranking result is quite different. For example, in the study of urban ecosystem service value, hydrological regulation accounts for the largest proportion in various ESVs(Hui, Wenwu et al. 2017, Tianhai, field et al. 2018). The second largest proportion is waste treatment and biodiversity maintenance (Hui, Wenwu et al. 2017, Tianhai, field et al. 2018).

The essence of ESV change is the area change of different ecosystems and the difference of equivalent factors in different ecosystems. Therefore, the trend of the value of individual ecosystem services in Xishuangbanna from 1996 to 2016 is consistent (Figure 1)

**Figure 1.**
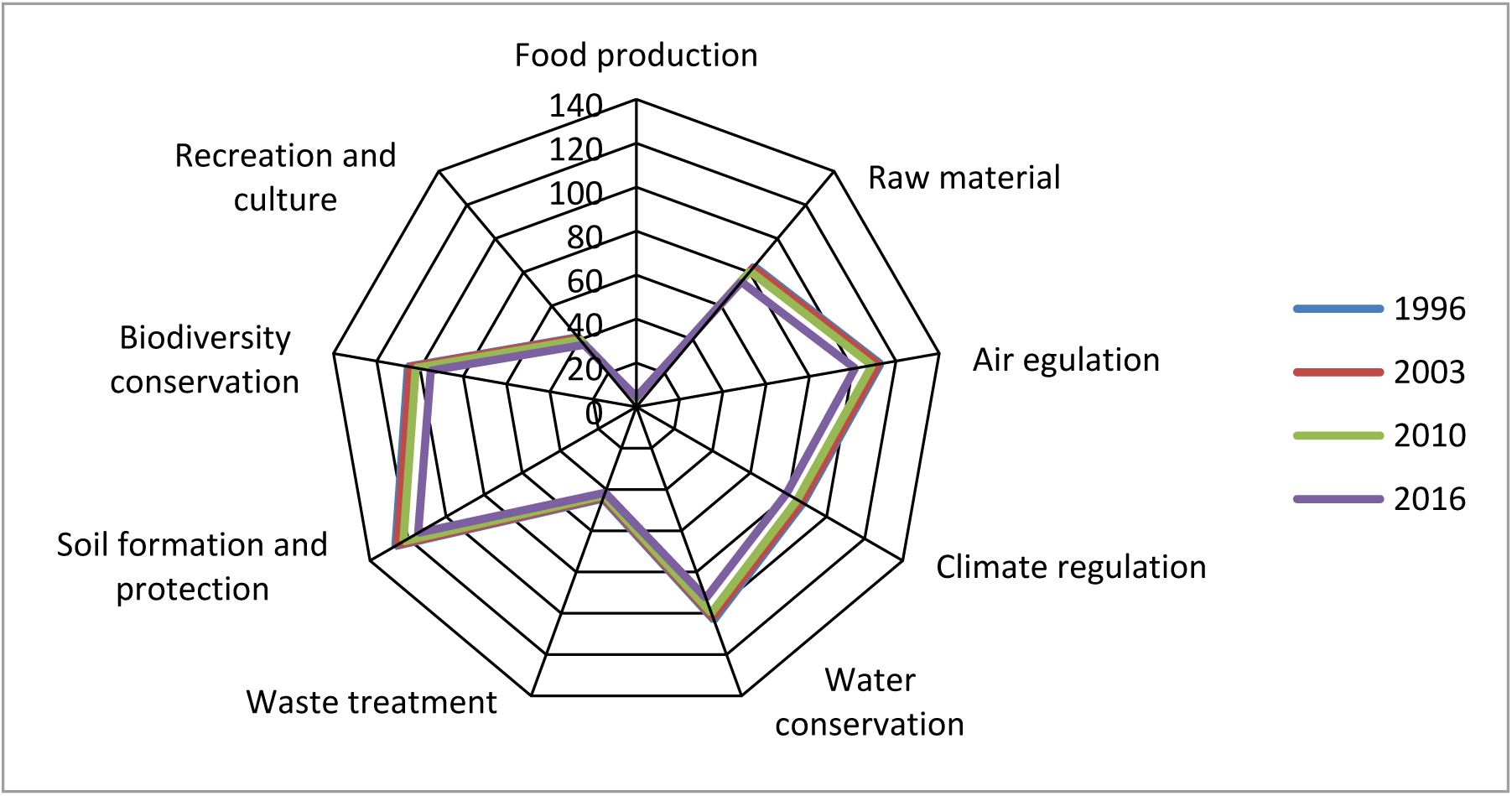
Structure of ecosystem service value in Xsihuangbanna, 1996-2016.

#### 2.2.2 ESV changes in the individual ecosystem services in Xishuangbanna from 1996 to 2016

According to Table 3, from the contribution of each component to the reduction of the total ESV, the contribution of ESV change in water conservation from 1996 to 2003 was the largest (17.29%). The contribution of ESV change to soil formation and protection from 2003 to 2010 was the largest (18.35%). The largest contribution of gas-regulated ESV changes during the period 2010-2016 (17.30%). The largest contribution of changes in soil formation and protection of ESV during the period 1996-2016 (17.17%).

**Table 3.** Ecosystem services values change and its contribution rate.

| Fisrt<br>classification | Second<br>classification | Value billion/ (亿元) |  |  |  | Contribution rate of value change/% |  |  |  |
| --- | --- | --- | --- | --- | --- | --- | --- | --- | --- |
|  |  | 1996 | 2003 | 2010 | 2016 | 1996-2003 | 2003-2010 | 2010-2016 | 1996-2016 |
| Supply services | Food production | 4.1512 | 3.9549 | 4.0058 | 4.4911 | 2.72 | -0.30 | -1.07 | -0.49 |
| Supply services | Raw material | 83.4826 | 82.6823 | 80.5464 | 74.2826 | 11.08 | 12.57 | 13.76 | 13.19 |
| Regulation services | Air regulation | 112.8175 | 111.8080 | 108.9442 | 101.0697 | 13.97 | 16.86 | 17.30 | 16.85 |
| Regulation services | Climate regulation | 87.5521 | 86.7146 | 84.5643 | 78.9300 | 11.59 | 12.66 | 12.38 | 12.36 |
| Regulation services | Water conservation | 103.2227 | 101.9738 | 99.8082 | 92.6174 | 17.29 | 12.75 | 15.80 | 15.21 |
| Regulation services | Waste treatment | 44.4557 | 43.7798 | 42.9431 | 41.5826 | 9.36 | 4.93 | 2.99 | 4.12 |
| Support services | Soil formation and protection | 126.6771 | 125.5667 | 122.4488 | 114.7029 | 15.37 | 18.36 | 17.02 | 17.17 |
| Support services | Biodiversity conservation | 105.6038 | 104.6699 | 101.9862 | 95.0425 | 12.93 | 15.80 | 15.25 | 15.15 |
| Cultural services | Recreation and culture | 41.3773 | 40.9656 | 39.8816 | 36.8908 | 5.70 | 6.38 | 6.57 | 6.43 |
| Total |  | 709.3401 | 702.1154 | 685.1285 | 639.6097 | 100.00 | 100.00 | 100.00 | 100.00 |

#### 2.2.3 Changes in the value of individual ecosystem services in different administrative regions of Xishuangbanna from 1996 to 2016

An analysis of the ecosystem service value of the three counties (Figure 2) shows that Xishuangbanna has higher ESV values in Erhai County and Mengla County. Xishuangbanna’s urban construction is dominated by Jinghong City (the seat of the state government), and tourism in Jinghong City is developing rapidly. Erhai County is not suitable for planting rubber, and the proportion of the tea garden ecosystem is relatively large. It is the main tea production place in Xishuangbanna.

**Figure 2.**
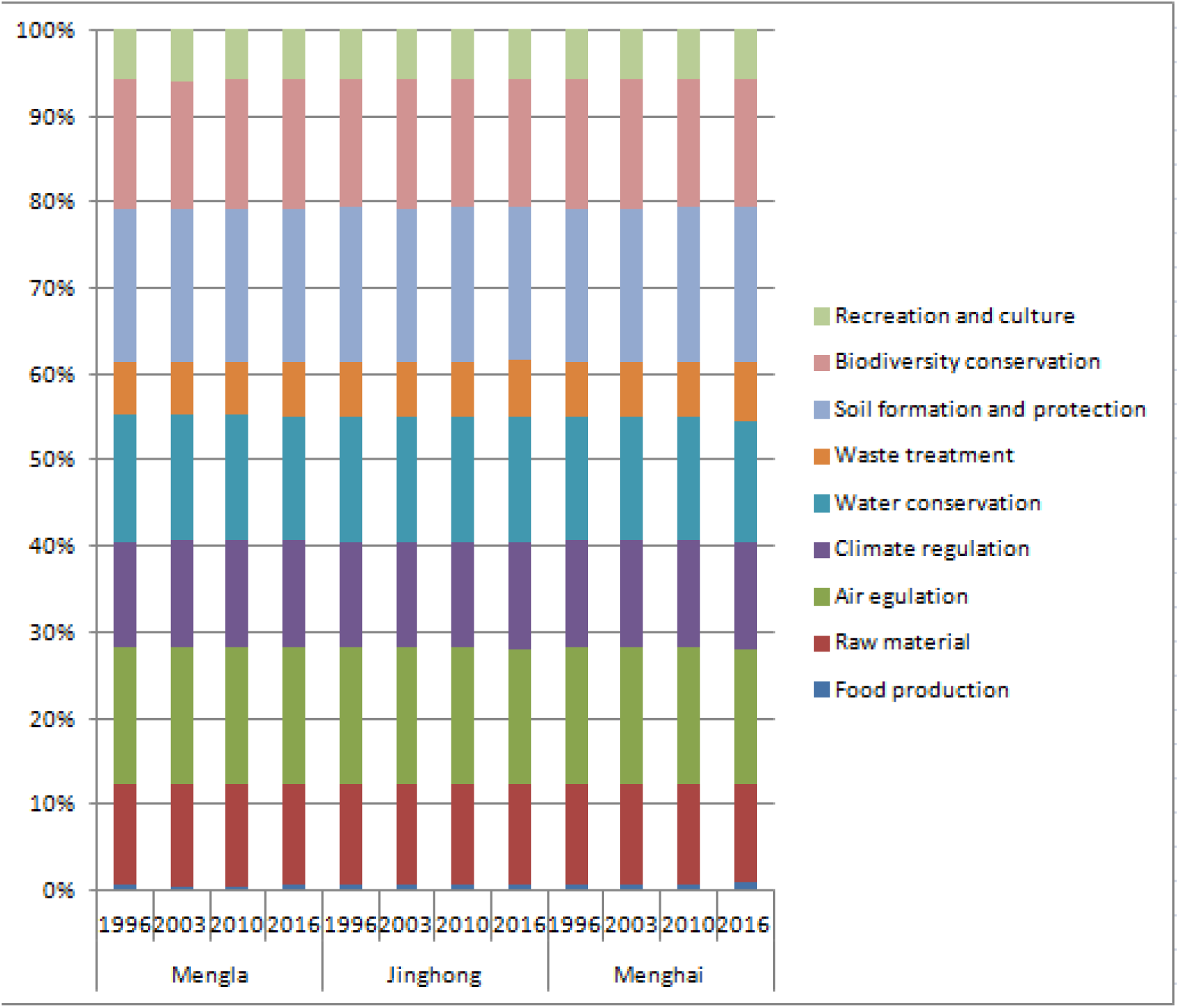
Changes in the value of individual ecosystem services in different administrative regions of Xishuangbanna from 1996 to 2016.

### 2.3 Spatiotemporal changes of ESV in Xishuangbanna

As can be seen from the figure 3, there are obvious spatial differences and temporal changes in ESV in Xishuangbanna. In 1996, land with an ESV in Xishuangbanna of more than 3,000 yuan / ha accounted for more than 90% of the total area. The ESV value was the highest in the year. ESV dropped in 2003, mainly at 1000-2000 yuan / ha. In 2010, ESV rose, but the area was larger than 3,000 yuan / ha in 1996, and the change was mainly concentrated in Erhai County. The area of ESV 1000-2000 yuan / ha in Erhai County increased. In 2016, ESV became more complicated and fragmented. High ESV areas were concentrated in Jinghong City and Mengla County, while Erhai County had fewer high ESV areas.

**Figure3.**
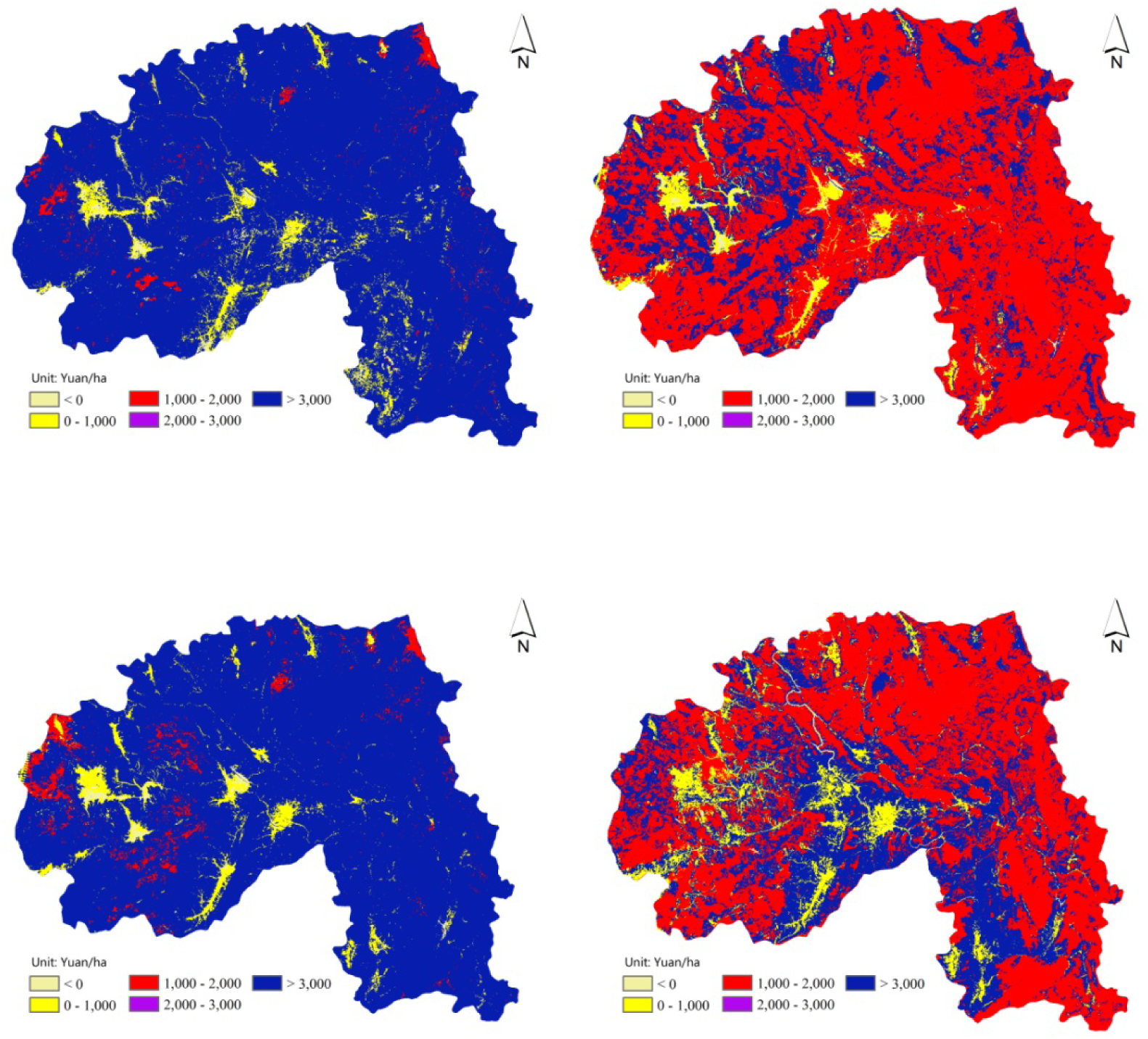
Spatiotemporal changes of ESV in Xishuangbanna from 1996 to 2016.

### 2.4 Ecosystem services sensitivity analyses

By calculation, the results of CS of all ecosystem types except forest in 1996, 2003, 2010 and 2016 were all lower than 1. The equivalence factors used in the study appear credible and accurate. This shows that the elasticity of the value coefficient of ESV is good, and the estimation results of this study are credible.

### 2.5 Cold and hot spot distribution ofESV in Xishuangbanna

Based on the Getis-Ord Gi * tool in the spatial analysis module, cold and hot spot identification was performed for the four ESVs of Xishuangbanna in 1996, 2003, 2010, and 2016, and the spatial distribution of the high and low ESV agglomerations in the fourth phase of the study area was obtained. Compared with 1996, the area of high-value ESV accumulation in Xishuangbanna in 2003, 2010 and 2016 has been significantly reduced. The hot spots of ESV in 1996 were mainly distributed in Jinghong City and Mengla County. Cold spots are distributed in Erhai County. Since 2003, 99% of cold spots have increased and hot spots have disappeared.

**Figure4.**
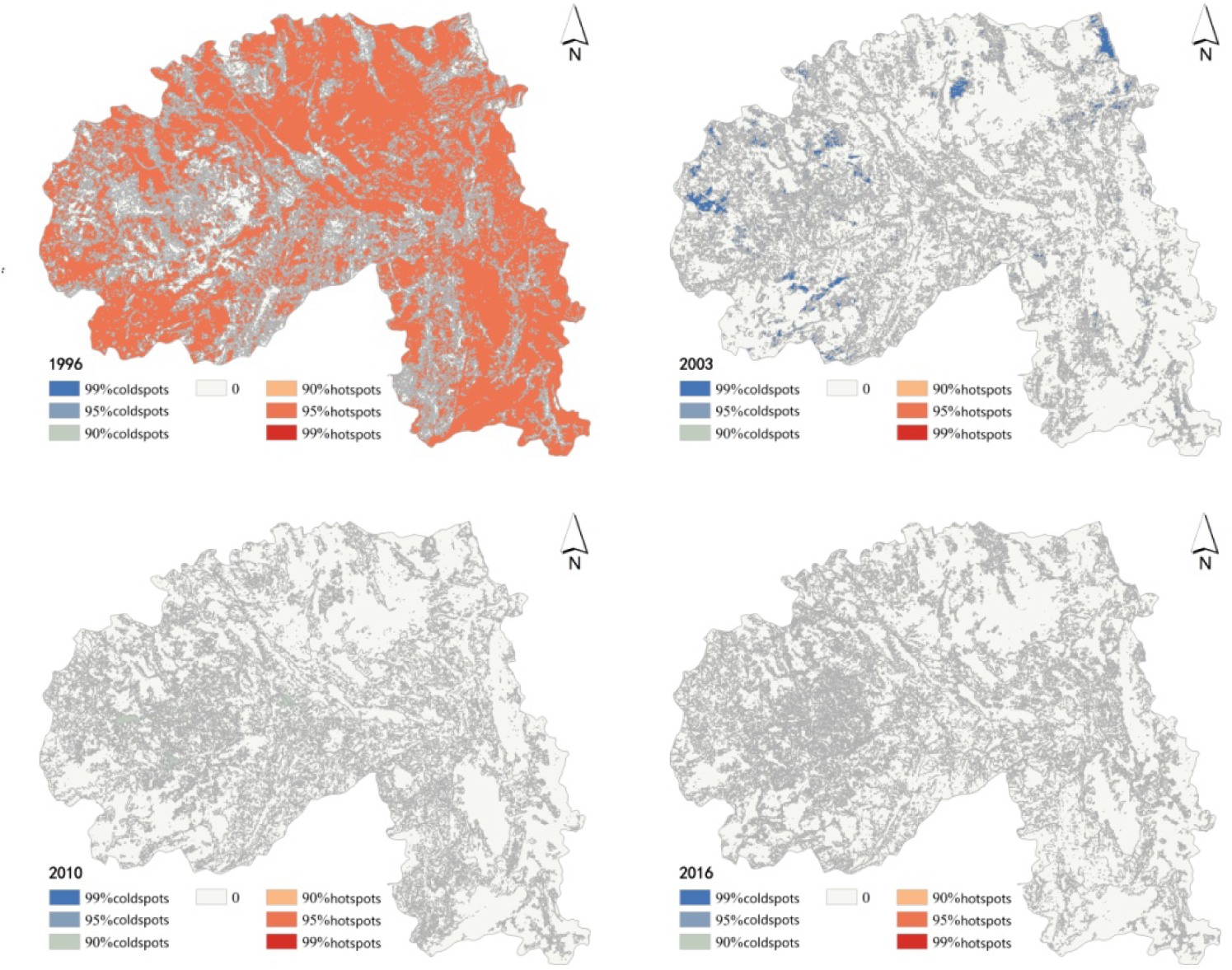
Cold and hot spot distribution ofESV in Xishuangbanna from 1996 to 2016.

## 3. Discussions and conclusions

1. When evaluating regional ESV, most scholars directly used the Xie Gaodi and other Chinese terrestrial ecosystems per unit area ecological service value equivalent scale(Gaodi, Chunxia et al. 2003, Xiaosai, Yongming et al. 2015), and used national scale equivalent factors to evaluate ESV in counties and cities, without reflecting regional differences.
2. During 1996-2016, the ESV in Xishuangbanna continued to decrease. In Xishuangbanna, due to the gradual decrease in the area of forest ecosystems, they have gradually been replaced by rubber, construction land, tea gardens, etc. in areas with low ecosystem service values. The overall value of ecosystem services in Xishuangbanna has continued to decline. As the forest land is the most important part of the ESV in Xishuangbanna, in the future development of Xishuangbanna, a corresponding ecological compensation system should be introduced to strengthen the protection of forest land, especially natural forests, and continue to promote the policy of “returning rubber to forests” to avoid excessive development and natural forest cover The replacement of economic forests leads to ecological imbalances and ecological risks, and eventually triggers international ecological security issues.
3. In this study, the ecological service value equivalent factor calculation method is used. Compared with other research results using the market value method, the calculation results are consistent, but the total value is smaller than the market value method(Zhaopeng and Youxin 2012) (Heli, Anyi et al. 2014).
4. There are obvious spatial differences and temporal changes in ESV and hot and cold spots in Xishuangbanna. In 1996, land with an ESV in Xishuangbanna of more than 3,000 yuan / ha accounted for more than 90% of the total area. ESV dropped in 2003, mainly at 1000-2000 yuan / ha. In 2010, ESV rose, and changes were mainly concentrated in Erhai County. The area of ESV 1000-2000 yuan / ha in Erhai County increased. In 2016, ESV became more complicated and fragmented. High ESV areas were concentrated in Jinghong City and Mengla County, while Erhai County had fewer high ESV areas. Compared with 1996, the area of high-value ESV accumulation in Xishuangbanna in 2003, 2010 and 2016 has been significantly reduced. The hot spots of ESV in 1996 were mainly distributed in Jinghong City and Mengla County. Cold spots are distributed in Erhai County. Since 2003, 99% of cold spots have increased and hot spots have disappeared.
5. The ESV assessment in this study is only for the first-class classification system, and there is no further subdivision of paddy fields and dry fields in cultivated land ecosystems, and there are woodlands and shrubs in woodland ecosystems.

## Acknowledgements

Our special thanks go to Prof Yang Yuming and Prof Ye Wen for providing constructive suggestions.

## Implications for conservation

### Funding

This work was supported by the 2017 Yunnan Provincial Department of Education Science Research Foundation Project (Grant 2017ZZX210).

